# Eliminating elevated p53 signaling fails to rescue skeletal muscle defects or extend survival in Lamin A/C-deficient mice

**DOI:** 10.1101/2022.07.08.499329

**Authors:** Tyler J. Kirby, Hind C. Zahr, Ern Hwei Hannah Fong, Jan Lammerding

## Abstract

Lamins A and C, encoded by the *LMNA* gene, are nuclear intermediate filaments that provide structural support to the nucleus and contribute to chromatin organization and transcriptional regulation. *LMNA* mutations cause muscular dystrophies, dilated cardiomyopathy, and other diseases. The mechanisms by which many *LMNA* mutations result in muscle-specific diseases have remained elusive, presenting a major hurdle in the development of effective treatments. Previous studies using striated muscle laminopathy mouse models found that cytoskeletal forces acting on mechanically fragile *Lmna*-mutant nuclei led to transient nuclear envelope rupture, extensive DNA damage, and activation of DNA damage response (DDR) pathways in skeletal muscle cells *in vitro* and *in vivo*. Furthermore, hearts of *Lmna* mutant mice have elevated activation of the tumor suppressor protein p53, a central regulator of DDR signaling. We hypothesized that elevated p53 activation could present a pathogenic mechanism in striated muscle laminopathies, and that eliminating p53 activation could improve muscle function and survival in laminopathy mouse models. Supporting a pathogenic function of p53 activation in muscle, stabilization of p53 was sufficient to reduce contractility and viability in wild-type muscle cells *in vitro*. Using three laminopathy models, we found that increased p53 activity in *Lmna*-mutant muscle cells primarily resulted from mechanically induced damage to the myonuclei, and not from altered transcriptional regulation due to loss of lamin A/C expression. However, global deletion of p53 in a severe muscle laminopathy model did not reduce the disease phenotype or increase survival, indicating that additional drivers of disease must contribute to the disease pathogenesis.

## Introduction

Mutations in the *LMNA* gene, which encodes the nuclear envelope proteins lamin A and C, cause a plethora of human diseases (‘laminopathies’) that include dilated cardiomyopathy, Emery-Dreifuss muscular dystrophy, *LMNA*-related congenital muscular dystrophy (L-CMD), and Hutchinson-Gilford progeria syndrome^1–5^. Although lamins A/C are nearly ubiquitously expressed, most mutations primarily affect skeletal and cardiac muscles^6^. The molecular mechanisms responsible for the striated muscle-specific defects remain incompletely understood. Although it is generally accepted that cardiac myopathy is the most significant contributing factor to early death in *Lmna*-mutant mouse models^7–9^, treatment approaches that specifically target skeletal muscle have been shown to improve longevity in laminopathy models^10^. Thus, further understanding the mechanisms that contribute to the skeletal muscle pathology in laminopathies could lead to more effective therapies.

We recently demonstrated that the mechanical forces in *LMNA* mutant skeletal muscle cells cause myonuclear damage, nuclear envelope rupture and DNA damage, and that the degree of DNA damage correlated with disease severity in both mouse models and human patients^11^. Nuclear envelope rupture can lead to DNA damage through multiple mechanisms, including mislocalization of DNA repair factors^12,13^ and exposure of DNA to cytoplasmic exonucleases^14^. Upon differentiation, skeletal muscle cells downregulate DNA repair factors^15^ and are less efficient at certain types of DNA repair compared to undifferentiated myoblasts^15,16^, leading to the possibility that skeletal muscle fibers are particularly susceptible to prolonged periods of DDR activation. Moreover, persistent activation of DDR pathways results in muscle fiber dysfunction during physiological aging^17^, further supporting the idea that elevated DDR signaling is detrimental to skeletal muscle function. Recent findings suggest that DNA damage and activation of DDR pathways could present a pathogenic mechanism in striated muscle laminopathies^12,18,19^. The tumor-suppressor protein p53 is one of the core factors that is activated in response to DNA damage^20,21^, and has recently been shown to be misregulated in striated muscle laminopathies ^18,19,22–25^. Under normal conditions, p53 is kept at low levels in the cell via the rapid targeting of p53 by E3 ubiquitin ligases, including MDM-2, Cop-1, Pirh-2 and ARF-BP1, that promotes its proteasomal degradation^26,27^. Upon activation of various DDR and stress response pathways, p53 is post-translationally modified, preventing its degradation and promoting p53 activation^21^. However, whereas the role of p53 in proliferative cells is well-established, with known involvement in cell cycle arrest, apoptosis, and senescence^28,29^, its role in terminally differentiated cells is still less well defined. This is despite the fact that in skeletal muscle, chronic activation of p53 can lead to muscle wasting^30^ and p53 has been implicated in other skeletal muscle diseases^31–33^. Thus, a better understanding of the functional consequences of p53 activation in striated muscle, and whether increased p53 signaling is a central driver of disease progression, could provide valuable insights into the mechanisms underlying muscle dysfunction in striated muscle laminopathies.

Here, we address the hypothesis that deregulation of p53 expression or increased p53 stabilization could result in skeletal muscle disease caused by *LMNA* mutations. A related question is the cause of the elevated p53 activation in striated muscle laminopathies. It is currently not known whether p53 activation in laminopathic striated muscle is the result of mechanically induced DNA damage or due to altered transcriptional regulation, as previously postulated for cardiac tissue^18,19^. If p53 activation is the result of mechanically induced DNA damage, we would expect that mutations that lead to reduced mechanical stability exhibited increased p53 stabilization and signaling, independent of altered p53 transcription. Furthermore, it is still unclear whether p53 activation is sufficient to induce muscle dysfunction and death in skeletal muscle cells, and if so, whether inhibiting p53 activation would improve skeletal muscle phenotypes in *Lmna* mutant mice. In the current work, we show that p53 stabilization and activity is elevated only in *Lmna*-mutant muscle cells that incur mechanically induced damage, and that mechanical damage to the nucleus is the primary driver of p53 stabilization, rather than altered transcriptional activity due to loss of lamin A/C expression. Furthermore, we demonstrate that chronic increase in p53 signaling is sufficient to induce muscle fiber dysfunction in wild-type muscle cells. However, global deletion of p53 in a severe striated muscle laminopathy mouse model did not improve the disease phenotype or increase survival, indicating that additional mechanisms must contribute to disease progression.

## Results

### In vitro differentiated Lmna mutant myotubes and myofibers have elevated p53 protein levels and transcriptional activity

To determine the expression and activity of p53 in *Lmna*-mutant myofibers, we differentiated primary myoblasts isolated from three different mouse models of striated muscle laminopathies, along with *Lmna* wild-type (*Lmna* WT) littermate controls, using an *in vitro* differentiation platform that allows for maintaining myofibers for 10 days (**Fig. 1A**)^34^. Lamin A/C-deficient mice (*Lmna*^-/-^, henceforth referred to as *Lmna* KO) develop severe muscular dystrophy and dilated cardiomyopathy and die around 6 to 10 weeks of age^7^. *Lmna* N195K (*Lmna*^N195K/N195K^)^8^ and *Lmna* H222P (*Lmna*^H222P/H222P^)^35^ mice are knock-in models based on human *LMNA* mutations. Both of these models exhibit muscular dystrophy and heart disease; however, whereas the *Lmna* N195K mice have an early disease onset and die around 10-14 weeks of age^8^, the *Lmna* H222P mice do not display a phenotype until around 12 (males) and 24 (females) weeks and die around 28 (males) and 40 (females) weeks of age^35^. Similar to the corresponding *in vivo* models, *in vitro* differentiated myofibers from these mouse models have varying degrees of ‘disease’ severity^11^. Myofibers derived from the more severe *Lmna* KO and *Lmna* N195K have compromised nuclear stability, extensive nuclear defects, including increased DNA damage, and reduced contractility and viability, whereas those from the later onset *Lmna* H222P model have normal myonuclear stability and show little nuclear abnormalities or functional defects^11^. Notably, despite differences in nuclear stiffness compared to wild-type cells, undifferentiated myoblasts from all three mouse models lack nuclear defects, presumably due to the limited cytoskeletal forces acting on the nuclei in these cells^11^.

**Figure 1.**
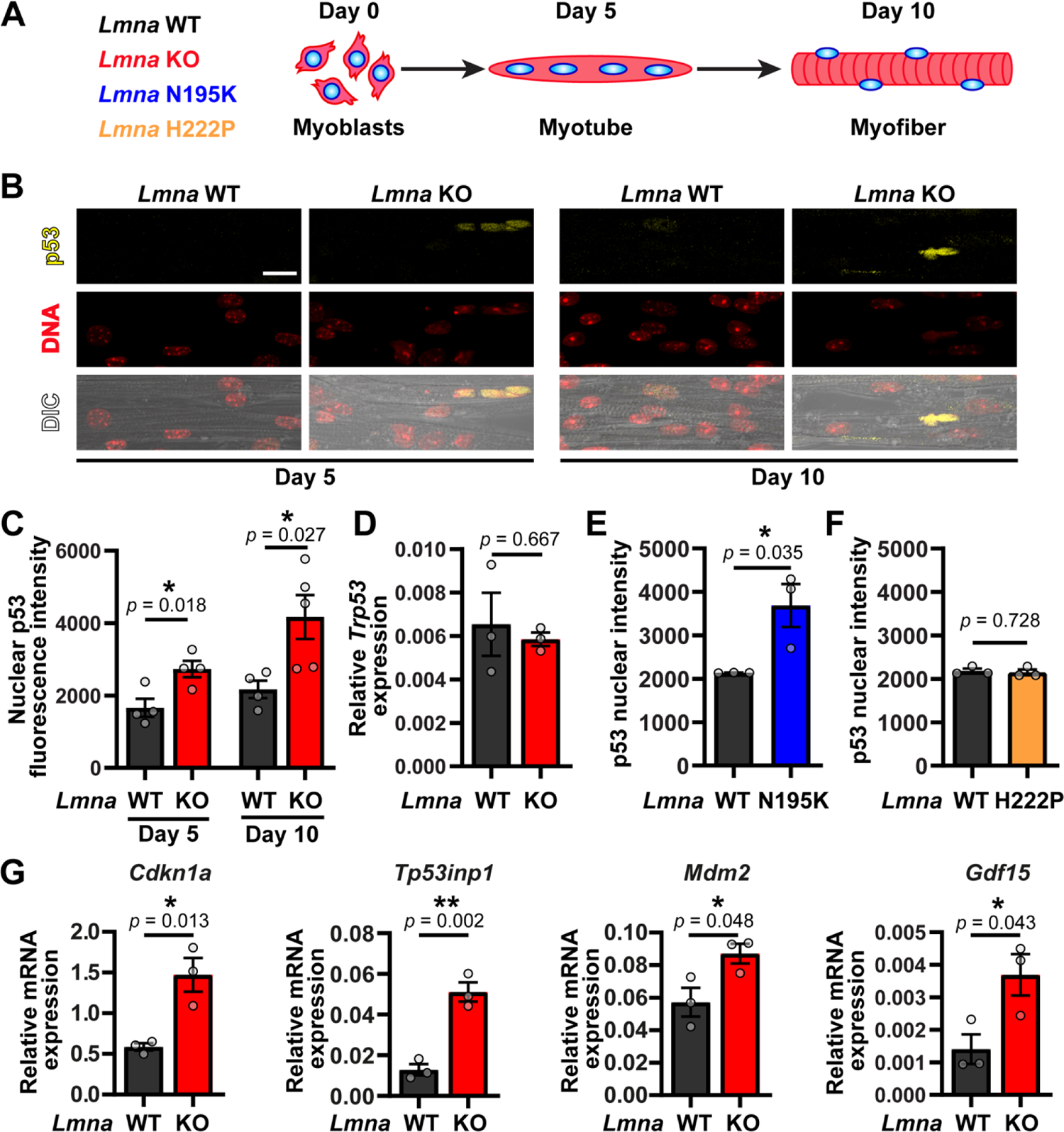
*Lmna* mutant myofibers display p53 stabilization and increased expression of p53-dependent genes. **(A)** Schematic of myoblast differentiation in the *in vitro* studies **(B)** Representative image of p53 immunofluorescence at day 5 and 10 of *in vitro* myofiber differentiation. Scale bar: 10 µm (**C**) Quantification of nuclear p53 immunofluorescence at day 5 and 10 of *in vitro* myofiber differentiation of *Lmna* WT and *Lmna* KO cells. (*N* = 4-5 independent cell lines with *n* = 113-420 nuclei quantified per genotype/time-point; Data shown as mean ± SEM; unpaired Student’s *t* test) (**D**) Quantification of relative gene expression of *Trp53* at day 5 of *in vitro* myofiber differentiation of *Lmna* WT and *Lmna* KO cells. (*N* = 3 independent cell lines; Data shown as mean ± SEM; unpaired Student’s *t* test) (**E**) Quantification of nuclear p53 immunofluorescence at day 10 of *in vitro* myofiber differentiation of *Lmna* WT and *Lmna* N195K cells. (*N* = 3 independent cell lines with *n* = 83-407 nuclei quantified per genotype/time-point; Data shown as mean ± SEM; unpaired Student’s *t* test) (**F**) Quantification of nuclear p53 immunofluorescence at day 10 of *in vitro* myofiber differentiation of *Lmna* WT and *Lmna* H222P cells. (*N* = 3 independent cell lines with *n* = 335-922 nuclei quantified per genotype/time-point; data shown as mean ± SEM; unpaired Student’s *t*-test) (**G**) Quantification of relative gene expression of p53-dependent genes (*Cdkn1a*, *Tp53inp1*, *Mdm2* and *Gdf15)* at day 5 of *in vitro* myofiber differentiation of *Lmna* WT and *Lmna* KO cells. (*N* = 3 independent cell lines; data shown as mean ± SEM; unpaired Student’s *t*-test)

Analysis of p53 protein levels in the *in vitro* differentiated myotubes and myofibers revealed results closely mirroring the time course and extent of nuclear damage in the different laminopathy models. Whereas nuclear p53 protein levels were indistinguishable between *Lmna* WT and KO undifferentiated myoblasts (Day 0) (**Suppl. Fig. 1 A-B**), *Lmna* KO myofibers had significantly elevated nuclear p53 protein levels at day 5 and day 10 of differentiation, compared to *Lmna* WT (**Fig. 1B and C**). These findings indicate that p53 protein levels are increased as the result of events that occur during differentiation, consistent with our previous findings of increased DNA damage at these time points^11^. Transcript levels for *Trp53*, the gene encoding p53, were not different at day 5 of differentiation (**Fig 1D**), suggesting that the increased p53 protein levels observed are the result of p53 stabilization, rather than increased mRNA expression. This finding is consistent with p53 protein levels being primarily controlled by post-translational modifications that significantly increase its turnover and half-life, which is extremely short (∼5-20 minutes) under homeostatic conditions^21^. Nuclear p53 protein levels were also significantly elevated in *Lmna* N195K myofibers at day 10 of differentiation (**Fig. 1E**), but not in *Lmna* H222P myofibers (**Fig. 1F**), suggesting that elevated nuclear p53 occurs in *Lmna* mutant cells with impaired nuclear stability and mechanically induced DNA damage. Finally, to determine whether the increased nuclear p53 was transcriptionally active, we performed qPCR for the canonical p53 target genes *Cdkn1a, Tp53inp1, Mdm2, and Gdf15*^36^ at day 5 of differentiation, i.e., the earliest time point at which we detected increased nuclear p53 levels in the *Lmna* KO myofibers. All of these target genes were upregulated in the *Lmna* KO myofibers (**Fig. 1G**), but not in undifferentiated *Lmna* KO myoblasts (**Suppl. Fig. 1C**), consistent with the nuclear p53 immunofluorescence results. These results demonstrate that loss of lamin A/C alone is not sufficient to result in activation of p53 and downstream genes. Collectively, our results demonstrate that p53 protein is stabilized specifically in *Lmna* mutant muscle cells that have reduced nuclear stability and that p53 stabilization occurs only following myoblast differentiation^11^.

### p53 transcriptional activity is also elevated in Lmna KO skeletal muscle fibers

To determine whether p53 activity was also altered in *Lmna* mutant mice *in vivo*, we examined p53 activity in skeletal muscle fibers isolated from *Lmna* WT and *Lmna* KO mice at 4 weeks of age. We focused on the myotendinous region of the muscle, as this region of the muscle fiber shows the greatest extent of nuclear and DNA damage^11^. Immunofluorescence staining revealed increased nuclear p53 levels in the *Lmna* KO myonuclei that appeared highly damaged (**Fig. 2A**). In contrast, we did not detect p53 in *Lmna* WT myonuclei (**Fig. 2A**). To confirm that p53 activity was elevated in *Lmna* KO skeletal muscle, we quantified the expression of p53 target genes, using the same genes that were upregulated in our *in vitro* differentiation system (**Fig. 1G**). Consistent with the *in vitro* data, *Lmna* KO skeletal muscle had increased expression of the p53 target genes *Cdkn1a*, *Tp53inp1*, *Mdm2*, and *Gdf14* (**Fig. 2B**). Interestingly, contrary to our findings in the *in vitro* differentiated fibers, *Trp53* transcript expression was also elevated in *Lmna* KO muscle (**Fig. 2B**), suggesting that additional mechanisms, or prolonged p53 activation, may lead to increased transcription of *Trp53* in *Lmna* KO skeletal muscle *in vivo*.

**Figure 2.**
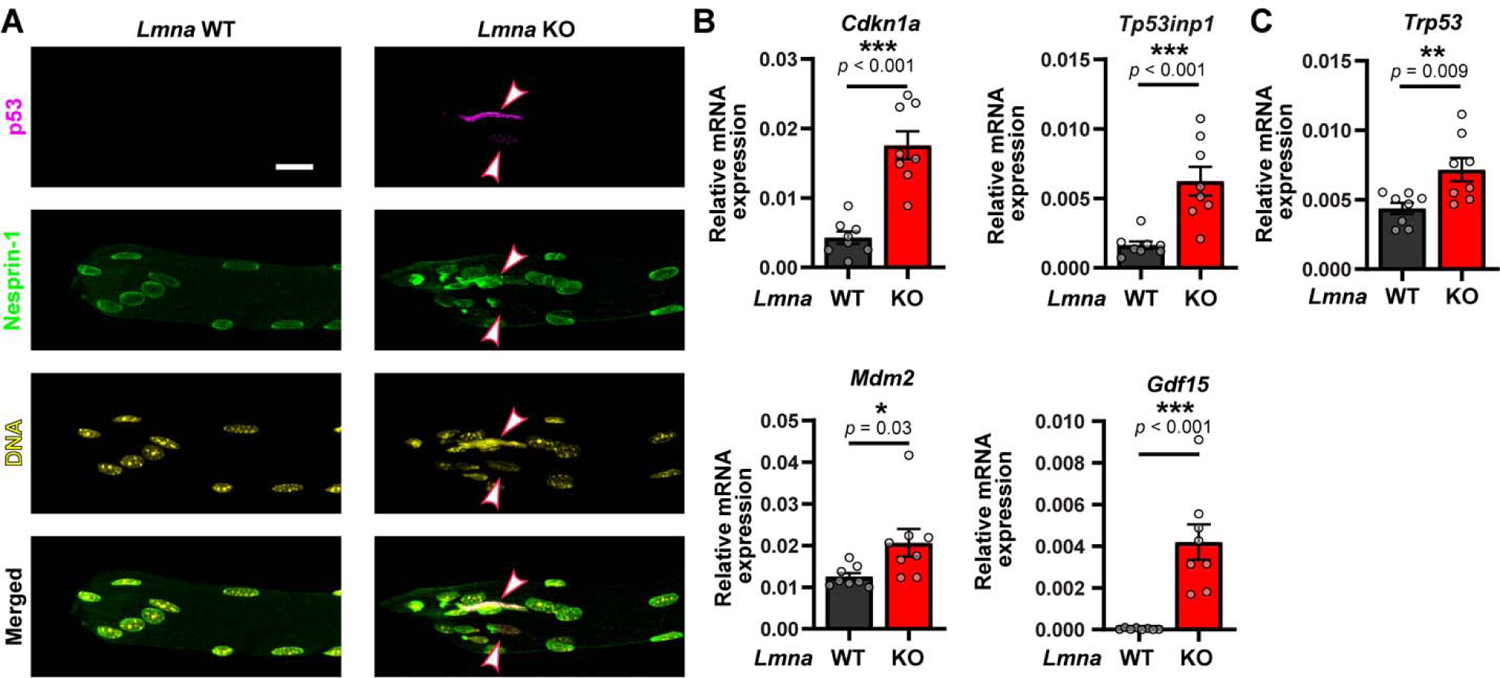
*Lmna* KO skeletal muscle displays increased expression of p53-dependent genes. **(A)** Representative image of p53 immunofluorescence in *Lmna* WT and *Lmna* KO skeletal myofibers isolated from the EDL. Note the increased levels in highly damaged myonuclei. Scale bar: 20 µm (**B**) Quantification of relative gene expression of p53-dependent genes (*Cdkn1a*, *Tp53inp1*, *Mdm2* and *Gdf15)* in *Lmna* WT and *Lmna* KO in the myotendinous region of the tibialis anterior muscle. (*N* = 8 mice; data shown as mean ± SEM; unpaired Student’s *t*-test) (**C**) Quantification of relative gene expression of *Trp53* in *Lmna* WT and *Lmna* KO skeletal muscle. (*N* = 8 mice; data shown as mean ± SEM; unpaired Student’s *t*-test)

### Increased p53 levels in Lmna KO myofibers is the result of mechanical damage to the nucleus

Two alternative, albeit not mutually exclusive, hypotheses could explain increased p53 activation in laminopathies affecting striated muscles. First, DNA damage resulting from mechanically induced damage to more fragile nuclei could lead to activation of DDR pathways, including p53, consistent with our previous report that preventing mechanical damage to the nucleus significantly reduces nuclear envelope rupture and DNA damage^11^, and our finding that increased p53 levels were only observed in differentiated myofibers, when increased cytoskeletal forces act on the myonuclei (**Fig. 1B and C, Suppl. Fig. S1A, B**). Alternatively, given the known role of lamins A/C in modulating chromatin organization and transcriptional regulation^37,38^, *Lmna* mutations may lead to altered gene expression that drives increased p53 activity in muscle cells^18,19^. To test these two hypotheses, we employed an approach where we disrupted the physical connection between the nucleus and cytoskeleton by expressing an inducible, dominant-negative construct of the KASH domain of nesprin-2 (DN-Kash2), which disrupts the Linker of Nucleoskeleton and Cytoskeleton (LINC) complex, thereby reducing the mechanical forces acting on the nucleus (**Fig. 3A-B**)^9,11,39^. A modified construct with two additional C-terminal amino acids, which also localizes to the nuclear envelope but does not interact with the inner-nuclear membrane SUN proteins (DN-Kash2Ext) and thus does not disrupt the LINC complex^40^, served as a negative control (**Fig. 3B**). In *Lmna* KO myofibers, disrupting the LINC complex at day 3 of differentiation reduced nuclear p53 levels to those of wild-type cells (**Fig. 3C, D**), whereas in *Lmna* WT myofibers, expression of the DN-Kash2 construct did not alter nuclear p53 levels relative to the control DN-Kash2Ext construct (**Fig. 3D**). These data indicate that the stabilization of p53 is primarily driven by force transmission to the nucleus, rather than simply due to the absence of lamin A/C.

**Figure 3.**
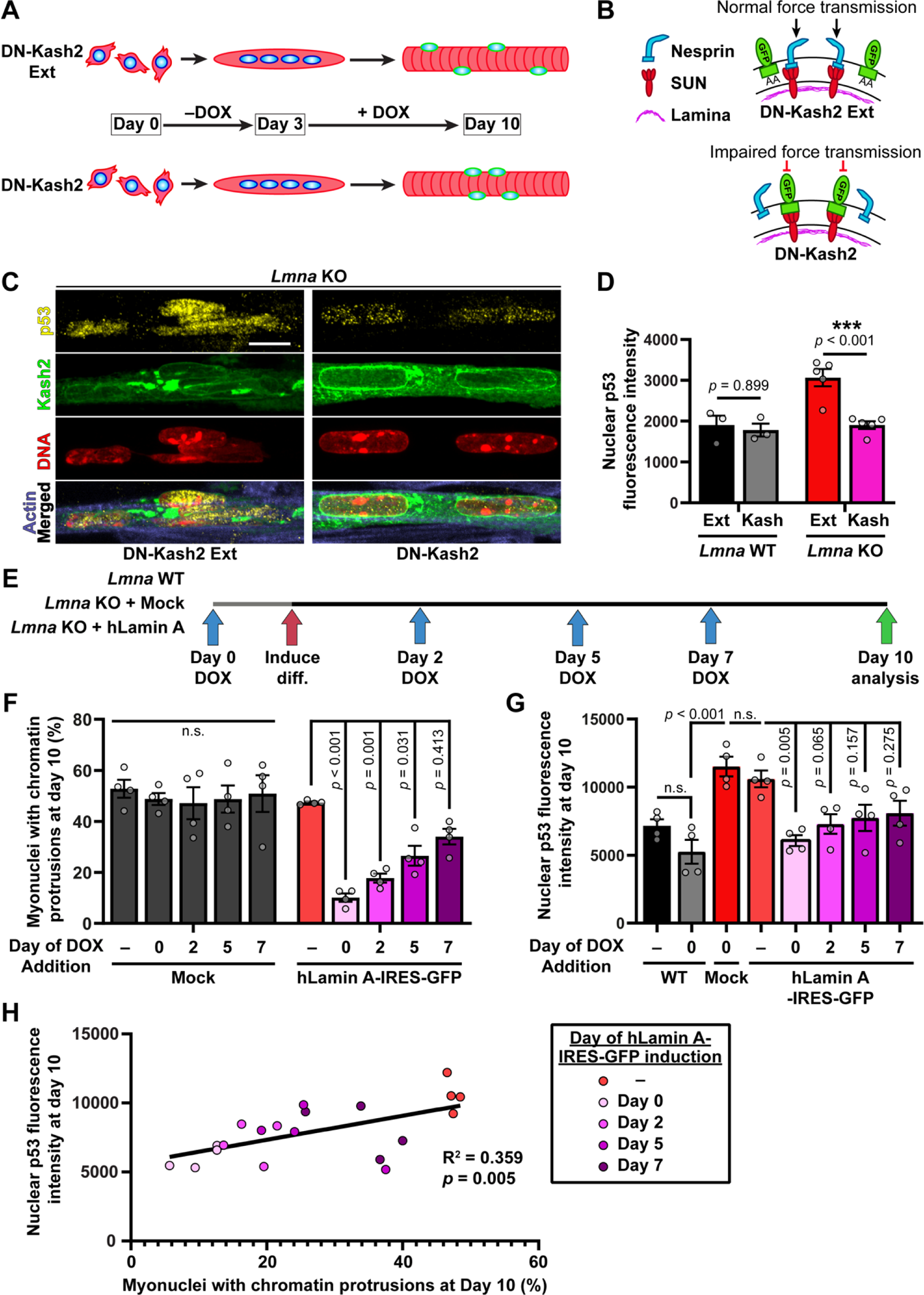
Increased nuclear p53 levels is the result of mechanical forces acting on nuclei in *Lmna* KO myofibers. **(A)** Schematic of LINC complex disruption studies, where the expression of either the DN-Kash2Ext (control) or DN-Kash2 construct is induced at day 3 of differentiation via the addition of doxycycline (DOX). (**B**) Schematic of LINC complex disruption by DN-Kash2 via preventing endogenous nesprins from interacting with the inner nuclear membrane SUN-domain containing proteins. The control construct, DN-Kash2Ext, contains an additional two alanine residues, which inhibits the interaction with SUN proteins, thereby maintaining the LINC complex. (**C**) Representative image of p53 immunofluorescence in *Lmna* KO myofibers at Day 10 of differentiation, expressing either the DN-Kash2Ext (control) or DN-Kash2 construct. Scale bar: 10 µm (**D**) Quantification of nuclear p53 immunofluorescence at day 10 of myofiber differentiation in *Lmna* KO skeletal myofibers expressing either DN-Kash2Ext (control) or DN-Kash2. (*N* = 3 (*Lmna* WT) or 5 (*Lmna* KO) independent cell lines with *n* = 95-533 nuclei quantified per genotype/time-point; data shown as mean ± SEM; two-way ANOVA with Sidak’s *post hoc* correction) (**E)** Schematic of experimental design for re-introduction of Lamin A in *Lmna* KO myofibers at various time-points during differentiation. Blue arrows indicate the different time-points when DOX was first added to the culture media (**F**) Quantification of chromatin protrusions in *Lmna* KO cells expressing Lamin A-IRES-copGFP or mock copGFP control at day 10 of differentiation. “Day” indicates the first day when expression was induced using doxycycline (DOX). (*N* = 4 (2 independent cell lines and 2 independent experiments) with *n* = 118-339 nuclei quantified per genotype/time-point. Data shown as mean ± SEM; Two-way ANOVA with Tukey’s *post hoc* correction) (**G**) Quantification of nuclear p53 immunofluorescence in *Lmna* WT or *Lmna* KO cells expressing either Lamin A-IRES-copGFP or mock copGFP control. “Day” indicates the first day when expression was induced using doxycycline (DOX). (*N* = 4 (two independent cell lines and two independent experiments) with *n* = 162-433 nuclei quantified per genotype/time-point; data shown as mean ± SEM; one-way ANOVA with Tukey’s *post hoc* correction) (**H**) Correlation between nuclear p53 levels and the extent of nuclear damage (percent chromatin protrusions) as a result of Lamin A-IRES-copGFP expression being induced at different timepoints during differentiation in *Lmna* KO muscle cells.

In a complementary approach, we tested whether re-introduction of lamin A into *Lmna* KO muscle cells at different stages of differentiation could prevent or reduce elevated p53 levels in these cells. If p53 activity was driven by altered transcriptional regulation due to the lack of lamin A, one would expect to see restoration of normal p53 levels regardless of when lamin A is re-introduced. On the other hand, if p53 activity was driven by mechanically induced nuclear damage, only introduction of lamin A during early stages of differentiation, i.e., when kinesin-1 mediated nuclear transport results in nuclear envelope rupture and DNA damage^11^, should reduce p53 activity. In contrast, later re-introduction of lamin A, i.e., after nuclear damage has already occurred, should be less effective in reducing p53 levels. To address this question, we generated cell lines expressing human Lamin A or mock control under the control of a doxycycline-inducible promoter. Doxycycline (DOX) was then added to the media at different time-points during differentiation (**Fig. 3E**). Re-expression of lamin A significantly reduced nuclear damage at all time-points, as evidenced by the fraction of *Lmna* KO myonuclei displaying chromatin protrusions, but the size of the effect strongly correlated with the timing of DOX administration (**Fig. 3F**). The largest rescue was observed when DOX was added prior to the onset of differentiation; the effect became increasingly less pronounced when lamin A was reintroduced later in differentiation, and re-expression of lamin A after 7 days of differentiation failed to significantly reduce nuclear abnormalities in *Lmna* KO myofibers (**Fig. 3F**). Analysis of nuclear p53 levels under the same conditions revealed a strikingly similar trend as observed for the myonuclear chromatin protrusions. A significant rescue of p53 levels back to *Lmna* WT levels occurred only in *Lmna* KO cells when ectopic Lamin-A was expressed at the earliest time-points (**Fig. 3G**), whereas later re-expression of lamin A failed to reduce nuclear p53 levels (**Fig. 3G**). These results indicate that re-expression of lamin A was not sufficient to reduce p53 levels if mechanically induced nuclear damage had already occurred. Further supporting the close connection between nuclear damage and p53 activation, we found a significant correlation between nuclear p53 levels and the extent of nuclear damage in *Lmna* KO myonuclei when analyzing data from all time points (**Fig. 3H**). Collectively, our results strongly suggest that mechanical damage to the nucleus is the primary driver of p53 stabilization and activation in *Lmna* KO skeletal muscle cells, and that preventing nuclear damage is sufficient to restore normal p53 levels, even in the absence of lamin A/C.

### Stabilization of p53 is sufficient to induce dysfunction in wild-type skeletal muscle cells

Although our results demonstrated that myonuclear p53 levels are elevated in striated muscle laminopathies that exhibit extensive nuclear damage in muscle cells, it remains unclear whether increased p53 activity directly affects muscle fiber function and viability. The effects of p53 activation in proliferating (i.e., cycling) cells has been well characterized, but functional consequences of p53 in post-mitotic cells, such as muscle cells, have yet to be fully explored. Furthermore, the majority of studies investigating p53 function in skeletal muscle have focused on loss-of-function of p53^41–44^, not stabilization and activation of p53. Thus, we set out to determine whether stabilization of p53 was sufficient to induce myofiber loss and/or dysfunction in wild-type muscle cells. To stabilize p53, we treated cells with nutlin-3, an inhibitor of the E3 ubiquitin-protein ligase Mdm2, which marks p53 for degradation, thereby resulting in increased p53 levels in nutlin-3 treated cells^45,46^. In our *in vitro* system, treatment with nutlin-3 for 24 hours resulted in a dose-dependent increase of nuclear p53 levels and a decrease in myofiber numbers, suggesting p53 induced cell death (**Fig. 4A**). For subsequent experiments, we selected a nutlin-3 dose that increased nuclear p53 levels, without inducing rapid cell death, thereby minimizing confounding factors. Wild-type *in vitro* differentiated myotubes were treated with 2.5 µM of nutlin-3, starting at day 5 of differentiation, and then analyzed for viability of myofibers at day 10 of differentiation (**Fig. 4B**). Nutlin-3-induced p53 stabilization resulted in a significant loss of myofiber viability compared to vehicle controls (DMSO) (**Fig. 4C**). The decrease in cell viability was the result of caspase-mediated cell death, as concurrent treatment with a caspase-3 inhibitor, Z-DEVD-FMK, restored viability to control levels (**Fig. 4C**). Furthermore, p53 stabilization reduced the contractility in the surviving myofibers (**Fig. 4D**), indicating that p53 activity reduces both myofiber viability and function.

**Figure 4.**
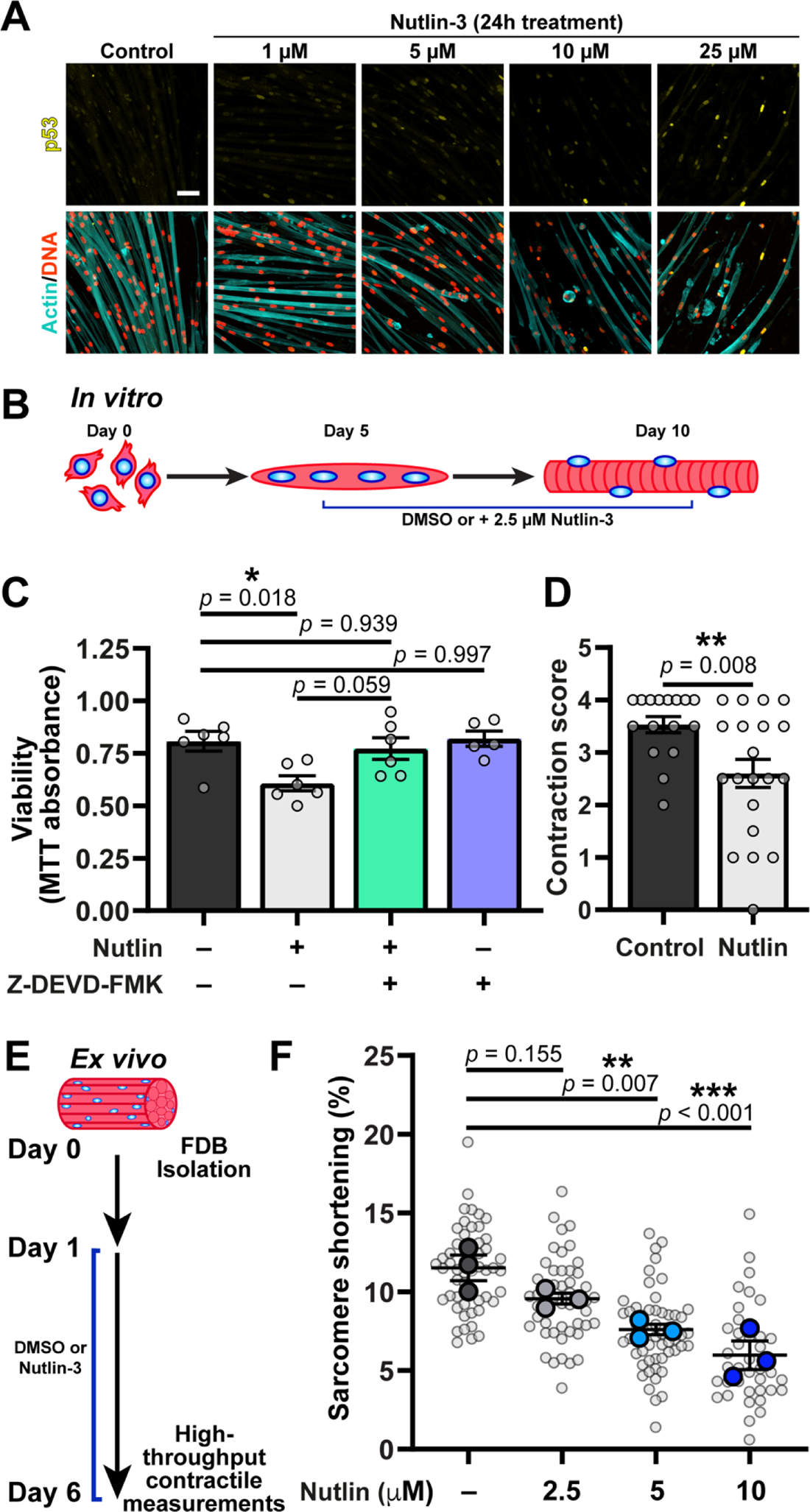
Stabilization of p53 is sufficient to induce myofiber dysfunction in *in vitro* differentiated and *ex vivo Lmna* WT myofibers. **(A)** Representative image of p53 nuclear immunofluorescence following 24 hours of treatment with increasing concentrations of nutlin-3 (MDM2 inhibitor). Note the decrease in cell viability with increasing concentrations of nutlin-3. Scale bar: 25 µm **(B)** Schematic of experimental design for nutlin-3 treatment in *in vitro* differentiated *Lmna* WT myofibers **(C**) Quantification of myofiber viability using the MTT assay. Cells were treated with 2.5 µM nutlin-3 and/or 50 µM Z-DEVD-FMK (caspase-3 inhibitor) starting at day 5 of differentiation. (*N* = 5 - 6 independent replicates from *n* = 3 independent cell lines; data shown as mean ± SEM; One-way ANOVA with Tukey’s *post hoc* correction (**D**) Quantification of myofiber contractility in *Lmna* WT myofibers following 5 days of 2.5 µM nutlin-3. (*N* = fields of view in *n* = 3 independent cell lines; data shown as mean ± SEM; unpaired Student’s *t* test) (**E**) Schematic of experimental design for nutlin-3 treatment in *ex vivo Lmna* WT muscle fibers. (**F**) Quantification of the percent sarcomere shortening in response to electrical stimulation in myofibers treated with different concentrations of nutlin-3 or DMSO control. (Data based on *N* = 3 mice and *n* = 38 – 55 muscle fibers; data shown as mean ± SEM; one-way ANOVA with Dunnett’s *post hoc* correction)

One limitation of *in vitro* differentiated myofibers is that even after 10 days of differentiation, they do not reach the maturity of muscle fibers *in vivo*, which may affect their contractility and response to elevated p53 levels. To overcome this limitation and to determine whether elevation of p53 affects contractility of mature muscle fibers, we performed contractility measurements on muscle fibers treated with nutlin-3 *ex vivo*. Muscle fibers were isolated from the flexor digitorum brevis (FDB) muscle of wild-type mice and then cultured *ex vivo* for 5 days in the presence of either nutlin-3 or vehicle control (**Fig. 4E**). Muscle fibers were then assessed for contractile function using a novel high-throughput system for assessing intact myofiber function (**Suppl. Fig. 2A**)^47^. Treatment with nutlin-3 significantly decreased fractional shortening (i.e., percent of contraction) of the muscle fibers in response to electrical stimulation, in a nutlin-3 dose-dependent manner (**Fig. 4F**). The higher doses of nutlin-3 also significantly reduced contraction speeds (**Suppl. Fig. 2B-E**). Collectively, these results demonstrate that prolonged stabilization of p53 is sufficient to induce contractile dysfunction in fully mature wild-type muscle fibers.

### Global deletion of p53 does not improve body weight, grip strength, muscle morphology, or viability in Lmna KO mice

To test whether targeting the elevated p53 levels in *Lmna* KO skeletal muscle cells could improve muscle function *in vitro* and *in vivo*, we generated *Lmna* WT and *Lmna* KO mice with either homozygous or heterozygous deletion of *Trp53* ^48^, or expressing wild-type *Trp53*. We first isolated and differentiated primary myoblasts from these mice and found that deletion of p53 improved viability in both *Lmna* WT and *Lmna* KO cells (**Suppl. Fig 3A**). The loss of p53 expression negatively affected the *in vitro* differentiation process (**Suppl. Fig 3B**)^49,50^, making the results difficult to interpret, since undifferentiated myoblasts exhibit little nuclear damage or DDR activation^11^. Thus, although our *in vitro* results supported the hypothesis that enhanced p53 activity leads to reduced vitality of *Lmna* KO muscle cells, we next examined the effect of p53 deletion in *Lmna* KO mice. We assessed body weight, grip strength, myofiber cross-sectional area, fibrosis and longevity in *Lmna* WT and *Lmna* KO mice that were wild-type, heterozygous or homozygous for a *Trp53*-null allele. Global deletion of *Trp53* is known to accelerate tumorigenesis, but this does not occur until three to six months of age in mice^48^, i.e., much later than the maximal life span of *Lmna* KO mice. Despite the positive effects of p53 deletion on myofiber viability *in vitro* (**Suppl. Fig. 3**), we found that neither deletion of one or both *Trp53* alleles improved body weight (**Fig. 5A, B**) nor grip strength (**Fig. 5C**). In addition, muscle quality, as assessed by average myofiber cross-sectional area (**Fig. 5D-E**) and percent of fibrotic area (**Fig. F-G**), was reduced in the *Lmna* KO mice compared to wild-type controls and not improved when *Trp53* was deleted. Finally, median survival (**Fig. 5H**) was not different in *Lmna* KO mice with deletion of one or both *Trp53* alleles compared to *Lmna* KO mice expressing two wild-type *Trp53* alleles. These results indicate that despite the increased activation of p53 in the skeletal muscle of *Lmna* KO mice, targeting p53 is insufficient to reduce the disease phenotype, likely because additional pathways are triggered by myonuclear damage that promote muscle cell dysfunction and death.

**Figure 5.**
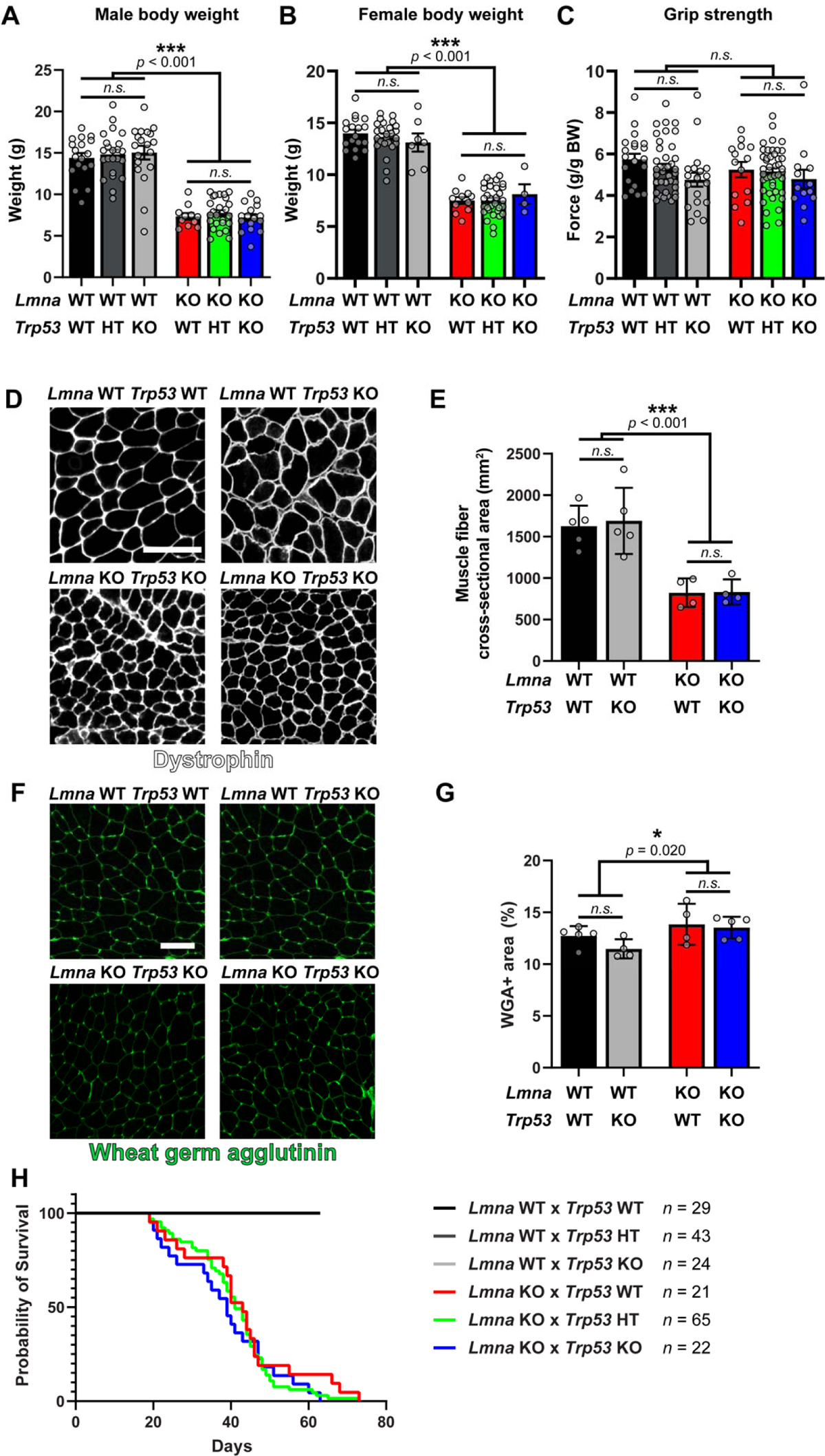
*Trp53* deletion does not improve muscle strength, body weight, muscle tissue morphology, or longevity in *Lmna* KO mice. **(A)** Body weight of 28 days old male *Lmna* WT and *Lmna* KO mice that are wild-type (WT), heterozygous (HT) or deficient (KO) for *Trp53*. (*N* = 11 - 25 mice; data shown as mean ± SEM; two-way ANOVA (*Lmna* genotype × *Trp53* genotype) with Tukey’s post-hoc correction) **(B)** Body weight of 28 days old female *Lmna* WT and *Lmna* KO mice that are wild-type (WT), heterozygous (HT) or deficient (KO) for *Trp53*. (*N* = 11 - 31 mice; data shown as mean ± SEM; two-way ANOVA (*Lmna* genotype × *Trp53* genotype) with Tukey’s *post hoc* correction) **(C)** Grip strength of 28 days old male and female *Lmna* WT and *Lmna* KO mice that are wild-type (WT), heterozygous (HT) or deficient (KO) for *Trp53*. (*N* = 13 - 43 mice; data shown as mean ± SEM; two-way ANOVA (*Lmna* genotype × *Trp53* genotype) with Tukey’s *post hoc* correction) (**D**) Representative image of tibialis anterior cross-sections from male *Lmna* WT and *Lmna* KO mice that are wild-type (WT) or deficient (KO) for *Trp53.* Tissue sections are immunofluorescently-labeled for dystrophin. Scale bar: 100 µm (**E**) Quantification of muscle fiber cross-sectional area (*N* = 4 - 5 mice; data shown as mean ± SEM; two-way ANOVA (*Lmna* genotype × *Trp53* genotype) with Tukey’s post hoc correction) (**F**) Representative image of tibialis anterior cross-sections from male *Lmna* WT and *Lmna* KO mice that are wild-type (WT) or deficient (KO) for *Trp53.* Tissue sections are labeled with wheat germ agglutinin (WGA) to stain for fibrosis. Scale bar: 50 µm (**G**) Quantification of muscle fibrosis area % calculated a percent of WGA area to total tissue area (*N* = 4 - 5 mice; data shown as mean ± SEM; two-way ANOVA (*Lmna* genotype × *Trp53* genotype) with Tukey’s post hoc correction) **(H)** Survival curves for male and female *Lmna* WT and *Lmna* KO mice that are wild-type (WT), heterozygous (HT) or deficient (KO) for *Trp53*. (*N* = number of mice)

## Discussion

The mechanisms by which *LMNA* mutations give rise to muscle-specific diseases have remained elusive, presenting a major hurdle in the development of effective treatments. We previously found striking correlations between the extent of nuclear abnormalities and DNA damage in skeletal myonuclei, scaling with the degree of disease severity in three mouse models of striated muscle laminopathies^11^. In addition, in skeletal muscle tissue biopsies obtained from laminopathy patients and healthy controls, the tissues from the most severe cases of muscular dystrophy had the highest levels of myonuclear DNA damage^11^. These findings led us to hypothesize that DNA damage and activation of DNA damage signaling might be pathogenic drivers of muscle disease in striated muscle laminopathies. We focused on p53, a key regulator of DDR and other cellular stress responses, because it has been shown that persistent activation of p53 leads to muscle wasting^30^ and that p53 is misregulated in laminopathic cardiac tissue^18,19,22^. The central role of the nuclear lamina in regulating chromatin localization and dynamics^37,51–53^ has led to the hypothesis that the pathogenicity of *Lmna* mutations is the result of an aberrant transcriptional profile in affected tissues. This mechanism is often referred to as the “gene regulation” hypothesis. The gene regulation hypothesis is supported by various studies that have demonstrated altered transcriptional patterns in *Lmna*-mutant cardiac and skeletal muscle cells and tissues, including increased p53 levels^19,54,55^. The extent to which altered gene expression is truly a driver of disease severity in laminopathy patients, however, is still not clear. Part of this uncertainty stems from the fact that transcriptional changes could occur downstream of other cellular events, including the nuclear damage often observed in cells with lamin A/C mutations, such as chromatin protrusions, nuclear envelope rupture, and DNA damage^11,56–59^. Our current data confirm that p53 is increased in the *Lmna*-mutant skeletal muscle fibers. Notably, however, the increase was mutation-specific and only observed in *Lmna*-mutant myofibers that have reduced nuclear stability (**Fig. 1**)^11^. Furthermore, increased p53 levels could not be rescued by reintroduction of lamin A after nuclei had already become damaged. In addition, we found that reducing mechanical damage to the nucleus by LINC complex disruption with a DN-Kash2 construct abrogated the increase in nuclear p53 in *Lmna* KO muscle cells (**Fig. 3A-D**). Collectively, these data indicate that p53 activation is the result of cellular events that are downstream of mechanical damage to the nucleus, rather than altered chromatin organization or transcriptional alterations caused by the loss of Lamin A/C.

One of the challenges of understanding disease mechanisms is distinguishing cellular events that are the primary cause of the disease from those that are secondary to disease progression. We previously showed that DNA damage and activation of DDR pathways occur in *Lmna* mutant myofibers that experience myonuclear damage and nuclear envelope rupture^11^; thus, we reasoned that the DDR could be a central driver of muscle dysfunction^17^. To this end, both our *in vitro* and *ex vivo* studies demonstrate that stabilization of p53 is sufficient to induce myofiber death and dysfunction (**Fig. 4, Suppl. Fig. 2**). How chronic stabilization of p53 negatively influences myofiber function is not clear, as the role of p53 in skeletal muscle has largely been investigated with loss-of-function studies^41,43,44,60^. Recent data suggests that in addition to its classically described role as a transcription factor, cytoplasmic p53 also plays important biological functions, including suppressing autophagy^61^, mitochondrial permeability and production of reactive oxygen species^62,63^, and modulation of Ca^2+^ homeostasis between mitochondria and endoplasmic reticulum^64^. Since our experiments were performed in intact myofibers, there are multiple mechanisms that could underlie the myofiber dysfunction, including impaired actin-myosin kinetics, improper calcium handling or mitochondrial dysfunction. Future studies are needed to determine the mechanism by which chronic p53 stabilization negatively affects muscle fiber function.

Surprisingly, despite the important role of p53 we identified for myofiber function and viability *in vitro*, deletion of *Trp53* was not sufficient to improve longevity or muscle function or morphology in *Lmna* KO mice (**Fig. 5**), unlike interventions such as LINC complex disruption that reduce mechanical stress on *Lmna* KO muscle nuclei and prevent nuclear damage^9,65^, or that prevent activation of the cGAS/STING pathway^66^, which can be triggered by exposure of genomic DNA to the cytoplasm following nuclear envelope rupture^67^. We hypothesize that part of the discrepancy between our *in vitro* and *in vivo* results can be attributed to differences in the ability for myofibers to undergo apoptosis. We previously reported increased caspase-mediated apoptosis of *Lmna* KO myofibers *in vitro*^11^, whereas muscle fiber loss *in vivo* is typically the result of necrotic mechanisms^68,69^. In addition, while we chose to focus on classic markers of muscle function in our *in vivo* studies, we cannot rule out that p53 deletion may affect other characteristics associated with muscle pathology, such as immune cell infiltration, centralized nuclei or mitochondrial abnormalities. Collectively, our findings suggest that p53 activation is not a primary driver of disease progression in *Lmna* KO mice, but instead p53 is activated downstream or in parallel to additional cellular stress responses resulting from nuclear damage in striated muscle cells, such as activation of the cytosolic DNA sensor cGAS^66^. These, yet to be identified pathways act in a p53-independent manner and thus still result in disease in *Lmna* KO mice deleted for *Trp53*.

One interesting finding from our study was that although we did not find differences in *Trp53* transcript expression between the *Lmna* WT and KO myofibers in the *in vitro* system, *Trp53* transcript levels were elevated in the *Lmna* KO mouse model. Possible explanations for this discrepancy could be the differences in the onset, duration and/or extent of DNA damage between the *in vitro* system and the *in vivo* mouse model. Previously, we showed that myonuclear damage and nuclear envelope rupture results from microtubule-mediated forces during myonuclear positioning^11^, with incidences of nuclear rupture dramatically increasing between day 2 and 3 of differentiation in the *in vitro* system^11^. Thus, in the *in vitro* system, the myonuclear and DNA damage was relatively acute, having only occurred ≈48 hours before assaying for p53 transcript and protein levels. Acute DNA damage primarily increases p53 levels through increased protein translation and stabilization of p53, without a significant change in mRNA levels^28,70^. In the *Lmna* KO mouse model, on the other hand, it is likely that the DDR signaling occurs over a longer period of time, compared to the *in vitro* system, and thus additional pathways or feedback mechanisms may lead to increased *Trp53* transcription^26,70,71^. Another explanation for the increased *Trp53* expression in the *Lmna* KO muscle tissue could be the differential expression of post-transcriptional regulators of *Trp53* transcript levels. Recently, it was shown that the heterogeneous nuclear ribonucleoprotein K (hnRNP K) can bind to the 5’ terminal region of p53 mRNA and influence mRNA levels^72^. Similarly, p53 contains a natural antisense transcript, designated Wrap53, that regulates endogenous p53 mRNA levels and further induction of p53 protein by targeting the 5′ untranslated region of p53 mRNA^73^. Finally, the 3’ UTR of p53 is the target of many different microRNAs, lncRNAs and RNA-binding proteins that could influence mRNA stability^70,74^; however, the extent to which the 3’ UTR influences *Trp53* mRNA half-life has recently been brought into question^75^. Nevertheless, differential expression of factors that post-transcriptionally influence p53 mRNA levels in *Lmna* KO skeletal muscle could be contributing to the increased p53 mRNA levels observed *in vivo*.

Ultimately, findings from this study highlight the need to uncouple the changes in gene expression that are due to altered chromatin organization or chromatin accessibility from gene expression changes that are downstream of mechanically induced damage to the nucleus. One limitation of our current study is that chromatin organization or accessibility was not assessed in response to reduced mechanical stress to the nucleus (i.e., LINC complex disruption). Thus, we cannot exclude that LINC complex disruption, in addition to preventing myonuclear damage, also alters chromatin dynamics/organization and gene expression, as mechanical stress to the nucleus can alter chromatin architecture^76^. Unraveling the interplay between chromatin organization, gene expression, and the role of specific nuclear envelope proteins is still an active area of investigation^77–81^. In addition, we opted to perform the LINC complex disruption studies in the *Lmna* KO cell line and not in the *Lmna* N195K or H222P mutant lines, as the *Lmna* KO line shows the most severe phenotype^11^, necessitating the most substantial improvement in function. If preventing mechanical damage to the nucleus is the primary mechanism contributing to improved function in the *Lmna* KO line, then we would expect that LINC complex disruption would rescue the defects caused by any mutation that results in loss of nuclear stability. It will be important for future studies to ascertain whether LINC complex disruption improves cellular and tissue-level function solely in mutations associated with reduced nuclear stability, or also applies to a broad spectrum of other *Lmna* mutations responsible for striated muscle laminopathies.

A limitation of our experiments using p53 stabilization in wild-type *in vitro* differentiated myoblasts and in isolated muscle fibers is that both of these studies relied on pharmacological stabilization of p53, independent of DNA damage, which may trigger additional signaling events. Furthermore, our studies in *Lmna*-mutant muscle cells were largely focused on the nuclear, transcriptionally active pool of p53; it is quite possible that p53 stabilization with nutlin-3 also led to an increase in cytoplasmic p53 or additional off-target effects, which could additionally contribute to negatively influencing muscle fiber contractility. Another important consideration is that we performed the *in vivo* p53 deletion experiments in an extremely severe laminopathy mouse model (*Lmna* KO), where the severe phenotype could mask any potential benefit of reducing p53 activation. Furthermore, we cannot rule out the possibility that other DNA damage signaling cascades that lie upstream of p53 or are p53-independent, such as DNA-PK or PARP1, are potentially contributing to disease progression. Nonetheless, our results are consistent with recent data showing only subtle improvements in mortality with p53 inactivation in a cardiac-specific laminopathy model^18^.

In conclusion, we propose that mechanical damage to myonuclei is a primary driver of severe disease pathogenesis caused by *LMNA* mutations. Mechanistically, it will be important to identify how reducing mechanical damage to the nucleus improves laminopathic striated muscle function, and whether mechanical damage to the nucleus is the initiating event in the pathogenesis caused by the lamin A/C mutations that directly affect myonuclear stability. Furthermore, our results highlight the importance of trying to unravel the contribution of mechanical and transcriptional events in the development of the diverse laminopathy phenotypes, with the hopes of developing more targeted and effective therapies for these diseases.

## Materials and Methods

### Animals

*Lmna* KO (*Lmna*^-/-^)^7^, *Lmna* H222P (*Lmna*^H222P/H222P^)^35^, and *Lmna* N195K (*Lmna*^N195K/N195K^)^8^ mice have been described previously. *Lmna*^+/–^, *Lmna*^H222P/+^, and *Lmna*^N195K/+^ mice were backcrossed at least seven generations into a C57-BL/6 line (Charles River Laboratories). For each mouse model, heterozygous mice were crossed to obtain homozygous mutants, heterozygous mice, and wild-type littermates. *Lmna* mutant mice were provided with gel diet (Nutri-Gel Diet, BioServe) supplement to improve hydration and metabolism upon onset of phenotypic decline. p53 (*Trp53*^-/-^)^48^ mice were purchased from Jackson Laboratory (Stock No: 002101) in a C57BL/6 background. All mice were bred, maintained, and euthanized according to IACUC approved protocols at Cornell University. For *ex vivo* contraction studies, post-mortem tissue was obtained from animals sacrificed for other approved research projects and/or breeding surplus of the VU University in accordance with the European Council Directive (2010/63/EU) by permission of the Animal Research Law of the Netherlands.

### *In vivo* survival studies and grip strength measurements

Only mice that survived to P14 were included in survival studies. Mice were euthanized if body weight loss exceeded 20% from their maximal weight for two consecutive days or 15% body weight loss compared to the previous day. In addition, mice that were visually lethargic, exhibited labored breathing, and/or had an abnormal rough coat were euthanized based on the recommendation of animal husbandry staff blinded for genotype. Body weight was determined at 4 weeks of age. Grip strength of 4-5–week old mice was tested using the Bioseb Grip Strength Test BIO-GS3 running the BIO-CIS software, with the test being performed by the same assessor for the entire study. The mouse was placed on the grid attachment and allowed to grip the grid with all four paws. The mouse was held by its tail and slowly pulled horizontally until it released the grid. This method was repeated 5-times consecutively per day for 3 days. The force exerted by the mouse was recorded every 15 milliseconds until 5 seconds, by which time it would have released the grid. The force applied to the grid just before it loses grip is recorded as the peak tension. The force measurements were plotted on a graph against time. For most of the measurements, the curve was smooth, increasing steadily as the force exerted increased, with a decrease as the mouse released the grid. An attempt was excluded if the curve showed an unusual spike in the otherwise smooth curve, and an additional attempt was made to replace it so that there were 5 measurements total per day. The maximum value of the force measurement for each attempt was used in our analysis. Force measurements were normalized to the average body weight of the mouse taken each day of grip strength testing.

### Myoblast isolation

Primary myoblasts were isolated and cultured as described previously^11^. Briefly, cells were harvested from *Lmna* KO, *Lmna* N195K, *Lmna* H222P, and wild-type littermates between 3-5 weeks for *Lmna* KO mice, 4-6 weeks for *Lmna* N195K, and 4-10 weeks for *Lmna* H222P mice. Myoblasts from wild-type littermates were harvested at the same time. Muscles of the lower hindlimb were isolated, cleaned of fat, nerve and excess fascia, and kept in HBSS on ice until all mice were harvested. The muscles were digested in 4 ml:1 g of tissue wet weight in a solution of 0.5% Collagenase II (Worthington Biochemicals), 1.2 U/ml Dispase (Worthington Biochemicals), 1.25 mM CaCl_2_ (Sigma) in HBSS/25 mM HEPES buffer. Digestion was carried out in a 37°C water bath for a total time of 60 minutes. When tissues were fully digested, the reaction was quenched using equal volumes of DMEM supplemented with 10% fetal bovine serum (FBS) and 1% penicillin/streptomycin (D10 media, Gibco). The cell suspension was strained through 70 and 40 μm filters (Greiner Bioscience) sequentially to remove undigested myotube fragments and tendon. The cell suspension was centrifuged at 800 × g for 5 minutes and washed with 8 ml of D10 media for a total of four times. Cells were then resuspended in primary myoblast growth media (PMGM; Hams F-10 (Gibco) supplemented with 20% horse serum and 1% penicillin/streptomycin and 1 µl/ml basic fibroblast growth factor (GoldBio)) and plated onto a 2% gelatin coated flasks. Cells were maintained in culture on gelatin coated flasks with media changes every other day. All experiments were carried out prior to passage 12. Each independent experiment was performed on a different set of *Lmna* mutant mice and wild-type littermates such that each independent experiment was sourced from a different animal to account for heterogeneity in phenotype.

### Myoblast differentiation

Myoblasts were differentiated according to a protocol modified from Pimentel et al.^34^ Wells were coated with growth factor reduced Matrigel (Corning) diluted 1:100 with IMDM with Glutamax (Gibco). Pre-cooled pipette tips were used to avoid premature polymerization. Matrigel was allowed to polymerize at 37°C for 1 hour and the excess solution was aspirated. Primary myoblasts were seeded at a density of 50,000 cells/cm^2^ in PMGM. Cells were allowed to attach for 24 hours before being switched to primary myoblast differentiation media (PMDM) composed of IMDM with Glutamax and 2% horse serum without antibiotics. This time-point was considered day 0. One day after the onset of differentiation, a top coat of 1:3 Matrigel:IMDM was added to the cells and allowed to incubate for 1 hour at 37°C. PMDM supplemented with 100 ng/ml agrin (R&D Systems) was added to the cells and henceforth replaced every second day.

### Plasmids generation and cell modification

To generate the DN-Kash2 and DN-Kash2Ext constructs, GFP-Kash2 and GFP-Kash2Ext were subcloned from previously published plasmids^40^ and inserted into an all-in-one doxycycline inducible backbone (pPB-tetO-MCS-EIF1α-rtTA-IRES-Neo)^82^. For rescue experiments, human lamin A was inserted into an all-in-one doxycycline inducible backbone (pPB-tetO-preLamin A-EIF1α-rtTA-IRES-Neo). For PiggyBac modifications, myoblasts were transfected with 1.75 μg of the PiggyBac plasmid and 0.75 μg of a Hyperactive Transposase using the Lipofectamine 3000 reagent according to the manufacture’s guidelines. Cells expressing the constructs were purified via fluorescence-activated cell sorting (FACS). For induction of expression, cells were treated with 1 μM doxycycline.

### Processing of tissue for cross-sectional tissue analysis

The tibialis anterior muscle was carefully excised from 6-week-old animals, coated with a thin layer of optimal cutting temperature compound (OCT compound, Tissue-Tek®, Sakura) and pinned at resting length on a piece of cork. The tissue was then rapidly frozen in liquid nitrogen-cooled isopentane and stored at –80 °C until further analyses. Tissue was cryo-sectioned at –20°C into 7 μm thin sections and mounted on a positively-charged glass microscope slide for immunofluorescence staining.

### Isolation of single muscle fibers for immunofluorescence

Single muscle fibers were harvested in a protocol adapted from Vogler et al.^83^. As previously described, fibers were isolated from the extensor digitorum longus (EDL) of male and female *Lmna* KO and wild-type littermates at 5-6 weeks of age. Briefly, the EDL was isolated from the mouse and placed directly into a 1 ml solution of F10 media with 40,000 U/ml of Collagenase I (Worthington Biochemicals). The tissue was digested for 15-40 minutes depending on muscle size in a 37°C water bath with agitation by inversion every 10 minutes. The reaction was quenched by transferring the digestion mixture to 4 ml of PMGM. Single fibers were hand-picked from the digested tissue using flame polished glass Pasteur pipettes. When necessary, the tissue was further dissociated by manual pipetting with a large-bore glass pipet. Fibers were washed once in fresh media prior to fixation with 4% paraformaldehyde (PFA) for 15 minutes at room temperature and subsequent IF staining.

### Isolation of muscle fibers for ex vivo culture

The flexor digitorum brevis (FDB) muscles were dissected in pre-warmed (37°C) and pH equilibrated (5% CO2) dissection media, comprised of Gibco-MEM, high glucose, pyruvate (Thermo Fisher, Germany), 10% heat inactivated FBS and 1% penicillin-streptomycin. The fourth lateral tendon and its myofibers were excluded from the muscle. FDB muscles were digested in 5 ml of sterilized 0.2% collagenase I dissolved in dissection media. Muscles were then incubated in a tissue culture incubator at 37°C and 5% CO2 for 90 minutes. After digestion, the muscles were carefully put back in 3ml dissection media without collagenase to recover for 30 minutes prior to trituration.

To release the individual myofibers, the tips of two p1000 pipette tips were cut off in different bore sizes and smoothened over a flame. Starting with the tip with the widest opening, the muscles were pipetted up and down until all fibers were released and in suspension. The tendons were carefully removed with a regular pipette tip. Myoblasts and remaining tissue were removed through gravity sedimentation. This was done by adding the cell suspension to 10ml dissection media and let it set for 20 minutes at 37°C and 5% CO2. This step was repeated by removing 10ml media, resuspending the cells and adding it to 10ml of fresh dissection media and then allowing cells to settle and pellet. Almost all media was removed from the pellet of FDB fibers. The fibers were put back into suspension by adding 1ml pre-warmed (37°C) and pH equilibrated (5% CO2) culture media, which contains Gibco-MEM, high glucose, pyruvate (Thermo Fisher, Germany), 20% Serum Replacement 2 (Sigma-Aldrich, USA), 1% horse serum and 1% penicillin-streptomycin. For each mouse, four wells of a 24-well plate (Corning, NY, USA) were precoated with laminin (40-80 μg/ml final concentration, Sigma-Aldrich, USA) and preloaded with 460µl equilibrated culture media. 140µl cell suspension was added to each well and cells were given two minutes to sink and adhere before moving to the incubator. Cells were cultured at 37°C and 5% CO2 and maintained by changing half of the media every 2/3 days by prewarmed and equilibrated culture media.

### FDB contractility studies

For *ex vivo* contractility measurements, FDBs were plated on a laminin-coated 24-well plate following isolation and treated with four different concentrations of Nutlin-3. At first, the cell medium was removed completely from the cells and changed it to 1 ml of tyrode medium (NaCL: 134 mM, Glucose: 11 Mm, KCL: 5 Mm, MgSO_4_*7H_2_0: 1.2 mM, NaH_2_PO_4_*2H_2_0: 1.2 mM, Hepes: 12 mM, Sodium Pyruvate: 5 mMm CaCl: 1 mM, pH 7.4 at 25°C). This was done in a stepwise fashion to prevent disturbing the fibers. First, 500 µl of medium was removed and replaced with 500 µl of tyrode medium. After a 2-minute incubation step, 500 µl of the medium was replaced by 500 µl of new tyrode media. The order that the different treatments were measured was randomized and the researcher who performed the measurements was blinded to the treatments. All contractility measurements were performed at 25°C using a MultiCell High Throughput System (IonOptix, Westwood, MA)^47^. Fibers were electronically stimulated using custom pacer with a pulse duration of 5.0 ms duration, 10 V and 1 Hz. Cells that were visibly responding to the electrical stimulation were used for measurements and the region used for contraction determination was kept consistent across all fibers. Ten contraction cycles were recorded for each muscle fiber and only cells that produced smooth contraction transients were used for further analysis. 20-25 single muscle fibers were measured per condition/replicate. The data was analyzed with the custom Cytocypher desktop program. In this program, measured traces are visualized with different colors based on whether a transient curve can be adequately fitted to the data based on pre-specified criteria. A “red” trace is rejected by the program, the “blue” trace is accepted by the program and the “grey” traces were contractions rejected during the image acquisition. Only cells that had transients that were accepted for at least 6 traces out of 10 contractions were included in the final data sets.

### Pharmacological treatments

For preliminary experiments, myoblasts were differentiated using the standard protocol and treated with pharmacological treatments at various timepoints. For p53 induction, myofibers were treated with nutlin-3 (Cayman Chemicals) (Table 1) starting at day 5 of differentiation for *in vitro* studies and 24 hours after isolation for *ex vivo* studies. To block caspase-dependent apoptosis, *in vitro* myofibers were treated with the caspase-3-specific inhibitor Z-DEVD-FMK (50 μM, Cayman Chemicals), starting at day 5 of differentiation.

**Table 1.** List of antibodies and chemicals.

| Reagent | Category | Vendor | Catalog Number |
| --- | --- | --- | --- |
| P53 (1C12) | Primary antibody | Cell Signaling | 2524 |
| Nesprin-1 | Primary antibody | Gift from D. Hodzic | NA |
| Dystrophin | Primary antibody | DSHB | MANDYS1(3B7) |
| Wheat Germ Agglutinin | Fluorescent probe | Sigma | L 4895 |
| Alexa Fluor™ 488<br>Phalloidin | Fluorescent probe | Invitrogen | A12379 |
| DAPI | Fluorescent probe | Invitrogen | D1306 |
| Donkey anti-Mouse IgG<br>(H+L), Alexa Fluor 488 | Secondary antibody | Invitrogen | A-21202 |
| Donkey anti-Mouse IgG<br>(H+L), Alexa Fluor 568 | Secondary antibody | Invitrogen | A10037 |
| Donkey anti-Rabbit IgG<br>(H+L), Alexa Fluor 488 | Secondary antibody | Invitrogen | A-21206 |
| Nutlin-3 | Chemical | Cayman Chemicals | 10004372 |
| Z-DEVD-FMK | Chemical | Cayman Chemicals | 14414 |

### Gene expression studies

Total RNA was extracted at Day 0 and day 5 of differentiation (*in vitro* studies) or from the myotendinous junction portion of the tibialis anterior of 4 week old *Lmna* WT and KO mice (*in vivo* studies) using TRIZOL/Chloroform, followed by column-based purification (Qiagen RNeasy Kit). cDNA was generated from 250-1000 ng of total RNA using an iScript™ cDNA Synthesis Kit (Bio-Rad Laboratories). Quantitative PCR for each candidate gene was performed using LightCycler 480 SYBR Green I Master (Roche) with the following cycle conditions: 95°C for 10 min, 50 cycles at 95°C for 15 s, 60°C for 30 s, and 72°C for 30 s, and a final cooling step of 4°C for 30 s. Primer sequences are shown in Table 2. All transcripts were normalized to the geometric mean of the housekeeping genes *18S*, *Gapdh* and *Rn7sk*.

**Table 2.**
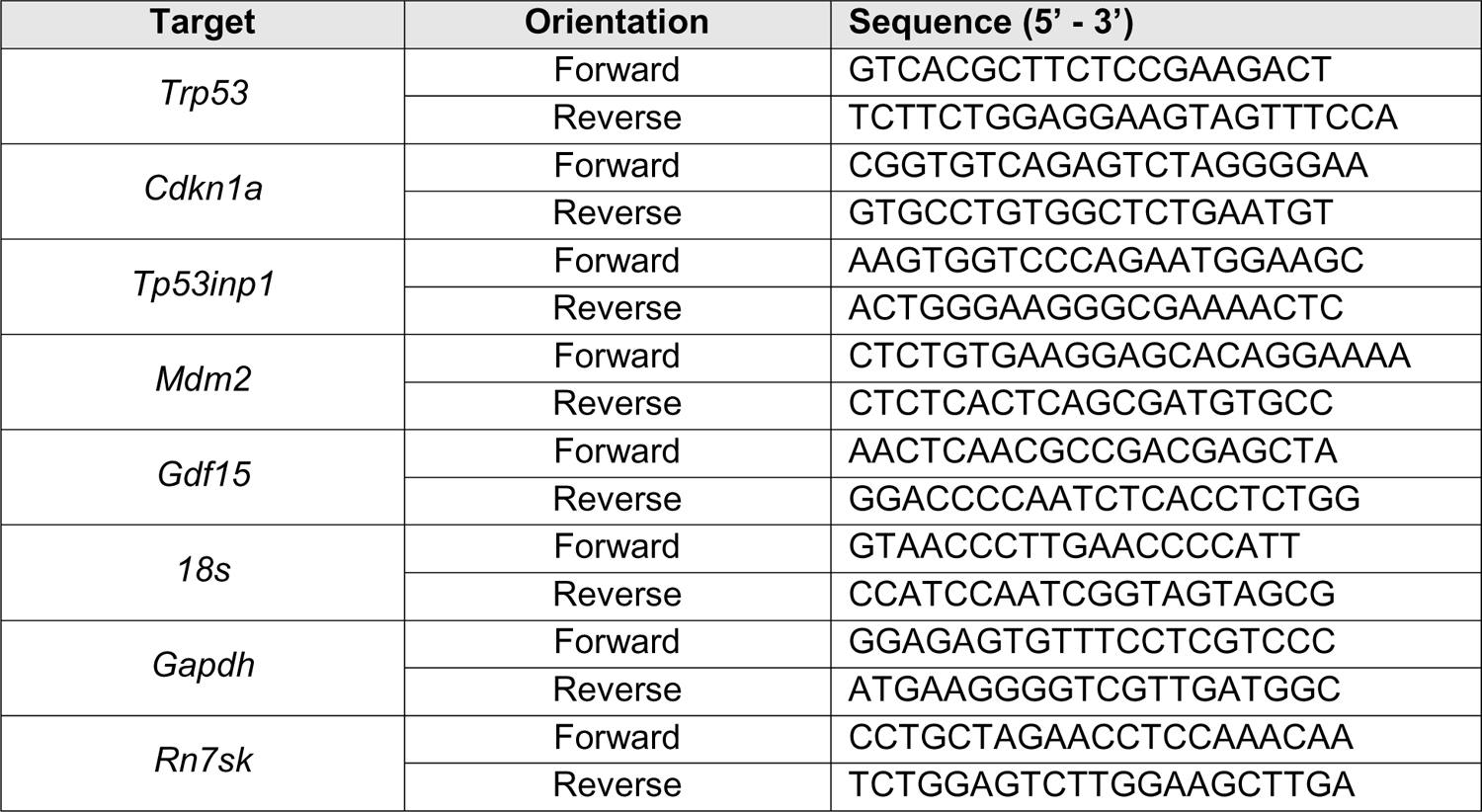
List of qPCR primers.

### MTT assay

Myoblasts, seeded in a 96-well plate and differentiated as previously described, were assayed for cell viability according to the manufacturer’s instructions (Promega, CellTiter 96 Non-Radioactive Cell Proliferation Assay). Fresh differentiation media was added two hours prior to the addition of 15 μl MTT 3-(4,5-dimethylthiazol-2-yl)-2,5-diphenyltetrazolium bromide dye. After incubation for 3 hours in MTT dye, 100 μl of Stop Solution was added to solubilize the formazan product (appears purple). Following overnight incubation at 37°C and 5% CO_2_, the absorbance of each well (measured at 590 nm) was analyzed using a microplate reader.

### Immunofluorescence staining of mouse cells and tissue

Cells were fixed in pre-warmed 4% PFA at the appropriate time point(s) and washed with PBS. Cells were blocked and permeabilized with a solution of 3% BSA, 0.1% Triton-X 100 and 0.1% Tween-20 (Sigma) for 1 hour at room temperature. Cells were stained with primary antibodies diluted in blocking solution according to Table 1 at 4°C overnight. Samples were washed with PBS and incubated for 1 hour at room temperature with 1:250 dilution of AlexaFluor antibodies (Invitrogen) and 1:1000 DAPI (Sigma). Single muscle fibers were stained using the same procedure in Eppendorf tube baskets with an increase in blocking solution Triton-X concentration to 0.25%. For tissue sections, Triton-X and Tween-20 were omitted from the blocking solution.

### Imaging acquisition

Cells on coverslips and mounted single muscle fibers were imaged with an inverted Zeiss LSM700 confocal microscope. Z-stacks were collected using 20× air (NA = 0.8), 40× water-immersion (NA = 1.2) and 63× oil-immersion (NA = 1.4) objectives. Airy units for all images were set between 1 and 1.5. Epi-fluorescence images were collected on a motorized inverted Zeiss Observer Z1 microscope equipped with CCD cameras (Photometrics CoolSNAP EZ or Photometrics CoolSNAP KINO) or a sCMOS camera (Hamamatsu Flash 4.0). Tissue sections stained with an anti-Dystrophin antibody for myofiber cross-sectional area analysis were imaged using the Aperio ImageScope (Leica Biosystems). Tissue sections stained with WGA for fibrosis analysis were imaged with an inverted Zeiss LSM900 confocal microscope using a 20× air objective (NA = 0.8) and the tiles function with support points.

### Image analysis

Image sequences were analyzed using ZEN (Zeiss) or ImageJ using only linear adjustments uniformly applied to the entire image region. Region of interest intensities were extracted using ZEN or ImageJ. For measurement of p53 nuclear intensities, projection images from z-stacks were generated using the “Sum Stack” function in ImageJ. A mask was generated from the maximum intensity projection of the DAPI-stained nuclear image and then applied to the p53 fluorescence image. For experiments that utilized a fluorescent reporter, a mask was first created using the fluorescent reporter channel to ensure only cells that were expressing the reporter at adequate levels were included in downstream analysis. Internal controls were utilized across independent experiments to normalize for differences in fluorescent intensity. Myofiber contractions were scored based on a minimum of 6 random fields of view per replicate using a blinded analysis as previously described^11^. Myofiber cross-sectional area was quantified using MyoVision^84^ where dystrophin was used to determine the cell boundary of the myofiber. For the WGA analysis, WGA-positive area was determined using manual segmentation by a researcher that was blinded to the genotypes. WGA-positive area was normalized to total tissue area to calculate the percent of tissue area stained positive for WGA.

### Statistical analysis

Unless otherwise noted, all experimental results were taken from at least three independent experiments or cell lines, and *in vivo* data were taken from at least three animals per group. We used either student’s t-tests (comparing two groups) or one-way ANOVA (for experiments with more than two groups) with post-hoc tests. When multiple comparisons were made, we adjusted the significance level using either Tukey or Dunnett corrections. Kaplan-Meier survival analysis was used to test for differences in survival rate. All tests were performed using GraphPad Prism. Unless otherwise noted, * denotes *p* ≤ 0.05, ** denotes *p* ≤ 0.01, and *** denotes *p* ≤ 0.001. Unless otherwise indicated, error bars represent the standard error of the mean (SEM). The data that support the findings of this study are available from the corresponding author upon reasonable request.

## Acknowledgements

The authors thank Colin Stewart for providing the *Lmna* KO and *Lmna* N195K mouse models, Gisèle Bonne for providing the *Lmna* H222P mouse model, Chloé Morsink and Sanna Luijcx for their help with the *ex vivo* contraction studies, and Alex Corbin for help with image analysis.

## Abbreviations

NE: Nuclear envelope

DNA: Deoxyribonucleic acid

p53: Tumor protein P53

WT: wild-type

LINC: Linker of nucleoskeleton and cytoskeleton

DOX: doxycycline

DNA-PK: DNA-dependent protein kinase

PARP1: Poly [ADP-ribose] polymerase 1

iPSC: induced pluripotent stem cell

FDB: flexor digitorum brevis

EDL: extensor digitorum longus

WGA: Wheat germ agglutinin

