## Supplemental Figures 1-3 for "Eliminating elevated p53 signaling fails to rescue skeletal muscle defects or extend survival in Lamin A/C-deficient mice" (from another version of this preprint: version 2, 2024-03-02)

**Supplementary figures**

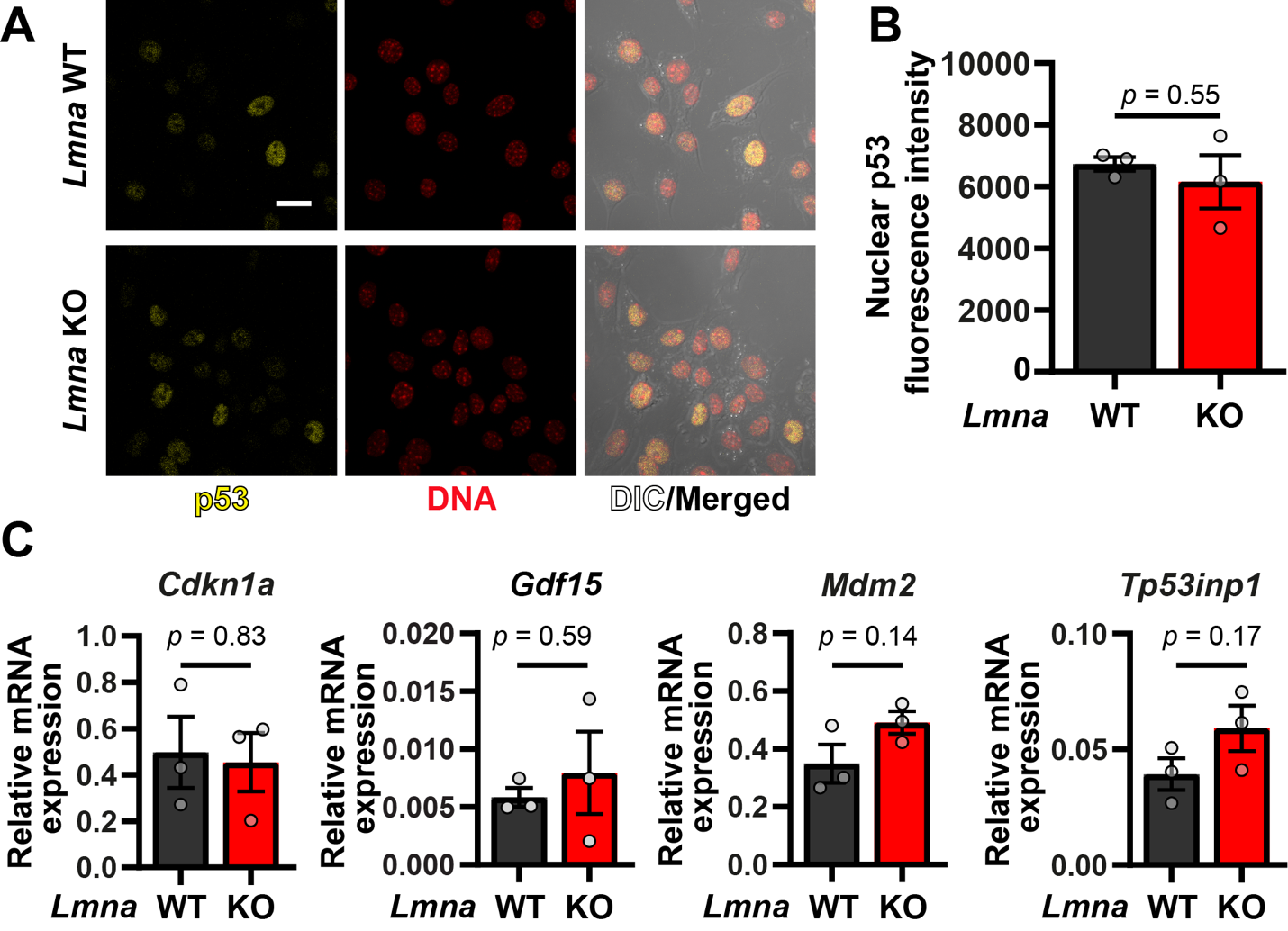

**Figure S1. p53 levels and p53-dependent genes are not differentially expressed in *Lmna* KO myoblasts prior to differentiation. (A)** Representative image of p53 immunofluorescence in primary myoblasts at day 0 of *in vitro* myofiber differentiation. Scale bar: 20 µM (**B**) Quantification of nuclear p53 immunofluorescence at day 0 of *in vitro* myofiber differentiation of *Lmna* WT and *Lmna* KO cells. *N* = 3 independent cell lines with *n* = 83-158 nuclei quantified per genotype. (**C**) Quantification of relative gene expression of p53-dependent genes (*Cdkn1a*, *Tp53inp1*, *Mdm2* and *Gdf15*) at day 0 of *in vitro* myofiber differentiation of *Lmna* WT and *Lmna* KO cells. *N* = 3 independent cell lines.

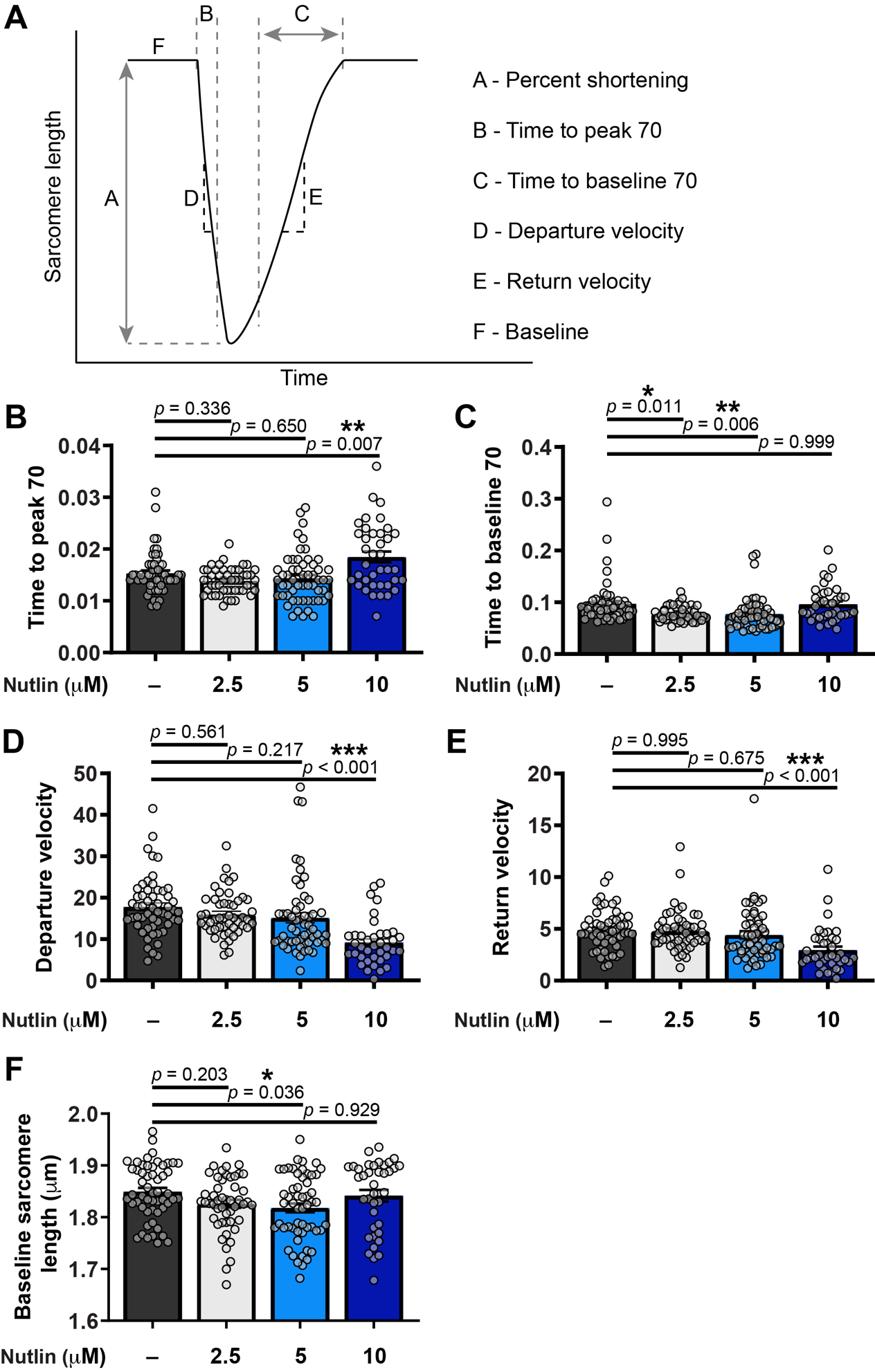

**Figure S2. p53 stabilization impairs muscle fiber contractility *ex vivo*. (A)** Representative contraction trace showing the contractile parameters measured using a high-throughput myocyte contraction system. **(B)** Quantification of the time required to reach 70% of peak sarcomere shortening (i.e. contraction time) in response to electrical stimulation in muscle treated with different concentrations of nutlin-3 or DMSO control. **(C)** Quantification of the time to return from 70% of peak sarcomere shortening back to baseline (i.e., relaxation time) in response to electrical stimulation in muscle treated with different concentrations of nutlin-3 or DMSO control. **(D)** Quantification of departure velocity (i.e., contraction velocity) in response to electrical stimulation in muscle treated with different concentrations of nutlin-3 or DMSO control. **(E)** Quantification of return velocity (i.e. relaxation velocity) in response to electrical stimulation in muscle treated with different concentrations of nutlin-3 or DMSO control. Data based on *N* = muscle fibers from *n* = 3 mice.

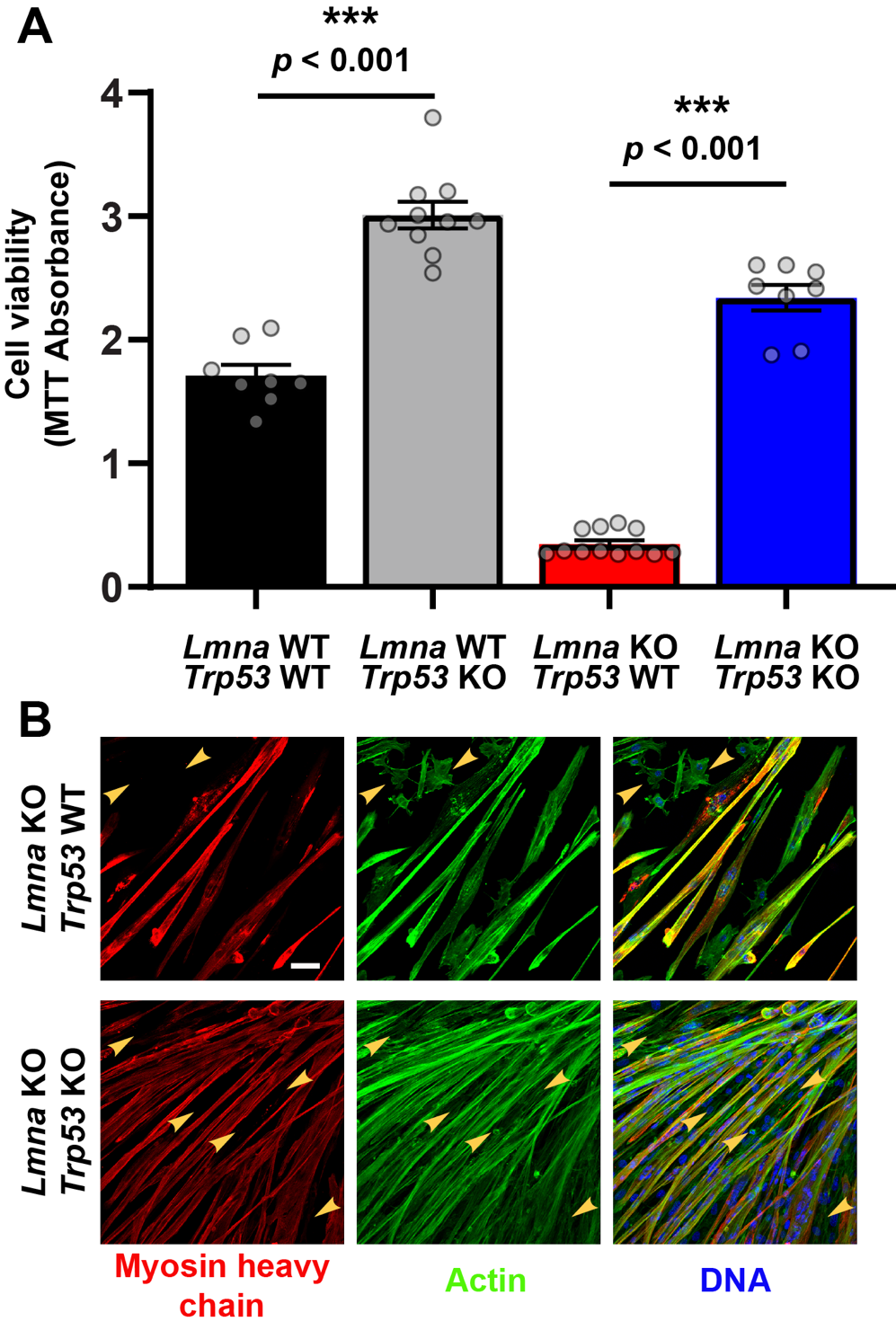

**Figure S3. Loss of p53 improves viability in *Lmna* KO muscle cells, but negatively affects myofiber differentiation. (A)** Quantification of myofiber viability using the MTT assay. *N* = independent replicates from *n* = 3 independent cell lines **(B)** Representative images of *Lmna* KO myofibers, either with (*Trp53* WT) or without p53 (*Trp53* KO) expression, at day 10 of differentiation. Arrows denote the presence of undifferentiated (Myosin heavy chain negative) cells, which increases with the loss of p53 expression. Scale bar: 50 µM.
